# Label-free monitoring of 3D cortical neuronal growth *in vitro* using optical diffraction tomography

**DOI:** 10.1101/2021.07.31.454602

**Authors:** Ariel J. Lee, DongJo Yoon, SeungYun Han, Herve Hugonnet, WeiSun Park, Je-Kyun Park, YoonKey Nam, YongKeun Park

## Abstract

The highly complex central nervous systems of mammals are often studied using three-dimensional (3D) *in vitro* primary neuronal cultures. A coupled confocal microscopy and immunofluorescence labeling are widely utilized for visualizing the 3D structures of neurons. However, this requires fixation of the neurons and is not suitable for monitoring an identical sample at multiple time points. Thus, we propose a label-free monitoring method for 3D neuronal growth based on refractive index tomograms obtained by optical diffraction tomography. The 3D morphology of the neurons was clearly visualized, and the developmental processes of neurite outgrowth in 3D spaces were analyzed for individual neurons.

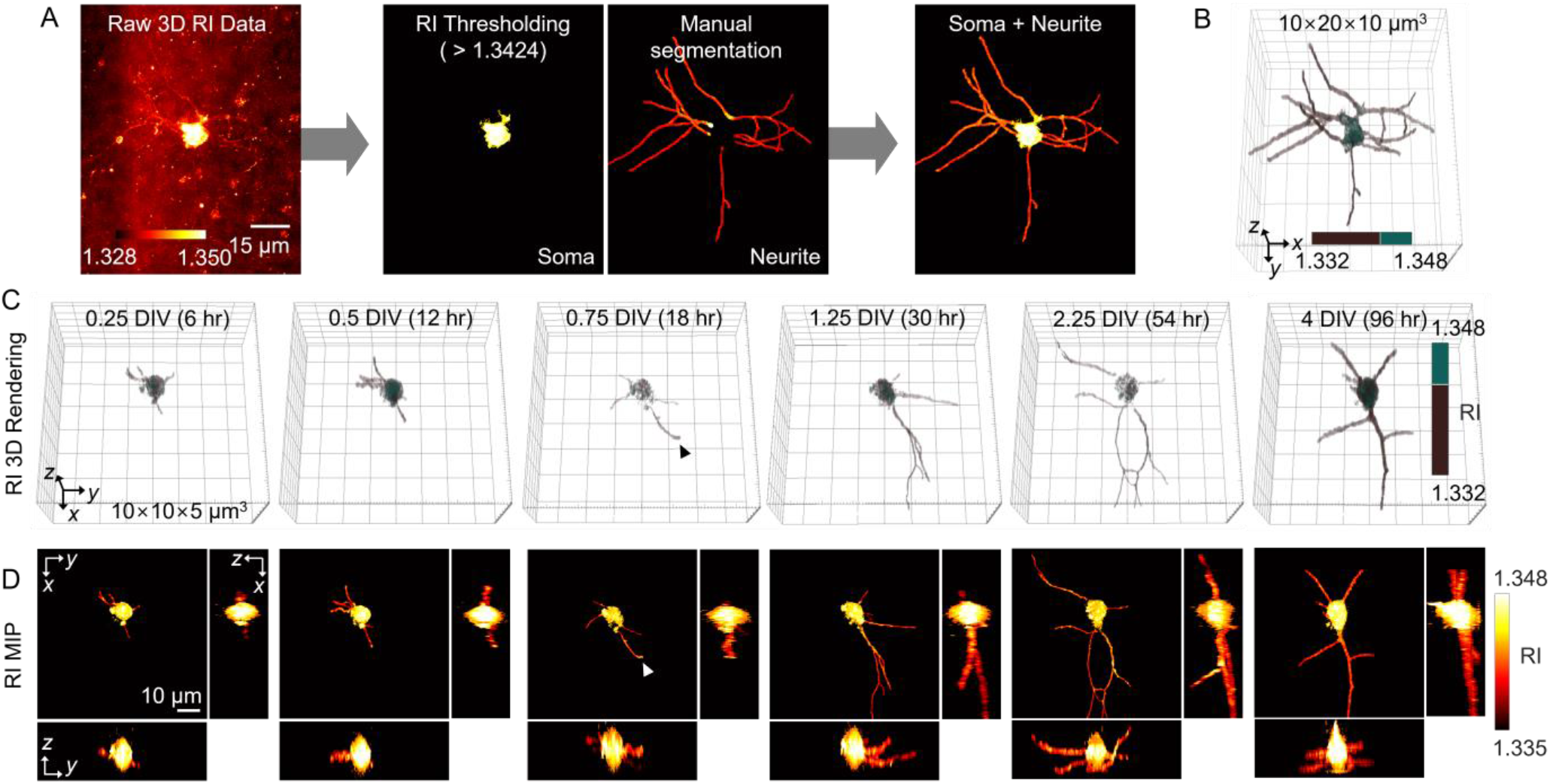

## 1. Introduction

The mammalian central nervous system has highly complex structures, which hinders the understanding of its pathophysiology. Culturing neurons derived from rats or mice *in vitro* serves as a platform for conducting experiments in a controlled environment [1, 2]. Established *in vitro* neuronal culture protocols adopt two-dimensional (2D) surfaces as substrates for neuronal growth [3–7]. *In vitro* primary neuronal culture systems have been widely utilized for investigating neuronal differentiation, polarity formation, network development, and maturation at both single- and multi-cell levels [8, 9].

Although numerous protocols have been developed for 2D neuronal cultures, the constraint in the culturing dimension significantly limits the ability of the culturing system to replicate *in vivo* microenvironments. The three-dimensional (3D) environment inherently provides a higher surface area for neuronal growth and allows outgrowth in all 3D directions with extended lengths [10]. Previous studies have shown that neurons cultured in a 3D environment exhibited significantly increased cell density [11], survival rate [12] and regenerative growth of axons [13], in addition to the electrophysiological activities that resembled those measured *in vivo* [14–16]. These findings have accelerated the development of numerous protocols [17, 18] and materials [19–23] to provide additional dimensions for the neurons to better mimic the 3D *in vivo* microenvironment. Matrigel, a natural extracellular matrix-based hydrogel, is widely used as a scaffold for 3D neuron cultures [10]. Cells embedded in Matrigel have been found to show a better survival rate and axon formation than 2D cultures [24–26]. 3D neural cell culture systems have been utilized for various regeneration studies [13, 27] and disease models [28].

These 3D culture platforms have revolutionized the *in vitro* culture to provide *in vivo-*like microenvironments for neurons. However, one of the major challenges in 3D culture platforms is the development of an imaging technique for volumetric visualization and analysis of cultured neurons. Volumetric visualization is essential for extracting useful information from 3D-structured neurons and analyzing their characteristics in full, as the morphology of the neurons has been found to affect their electrophysiological properties [29].

One of the most widely utilized solutions for this problem is fluorescence imaging based on laser scanning confocal microscopy [30, 31]. Conventional protocols require fluorescent labeling using immunofluorescence staining [32]. Although these protocols enable clear subcellular positioning of the labels with high specificity and can stain multiple targets, they require the fixation of a sample, which terminates any biochemical reaction in the sample. Thus, multiple samples must be prepared to study morphological changes during the development processes (Fig. 1). Optimizing the choice of chemicals and protocols is critical for successful imaging results [33]. As the cultured neurons undergo the fixation procedure for fluorescence imaging, multiple samples should be prepared to study neuronal growth at multiple time points. This implies that another set of resources is required for culturing and imaging to measure neurons for an additional time point. In addition, the sample preparation procedure, including fixation and staining, requires 4–5 days, which is labor-intensive and time-consuming [34]. Expressing fluorescent proteins by transfecting genes is another approach for labeling specific molecules for imaging [35]. However, this requires genetic modification, and it may cause unexpected effects on target proteins.

**Fig. 1.**
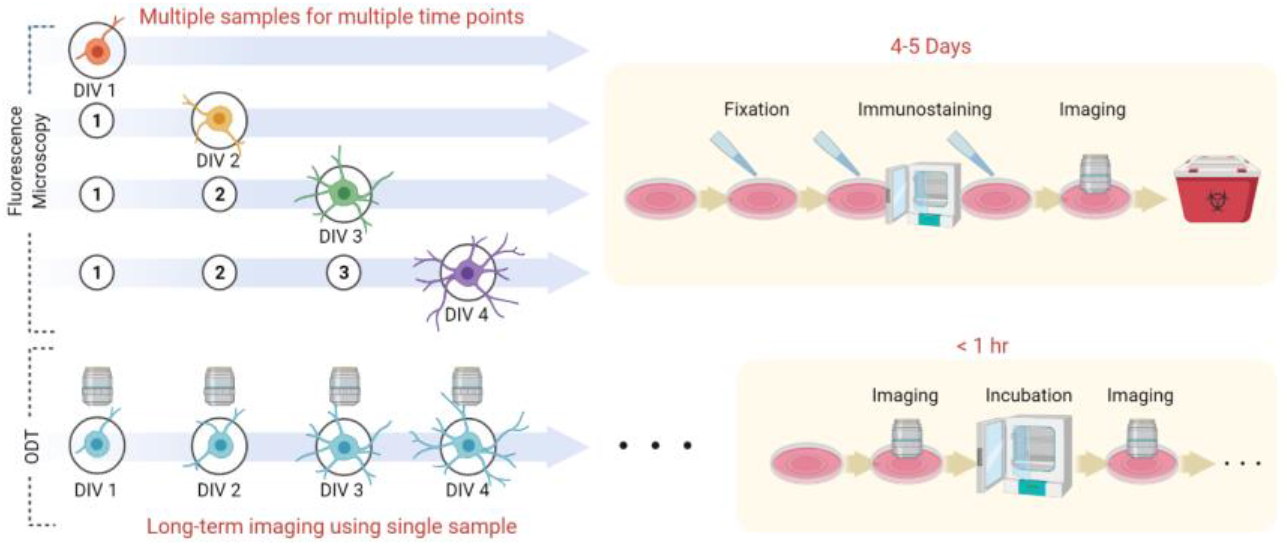
Advantages in experiment procedure for neuron time-lapse imaging in 3D when using optical diffraction tomography (ODT), compared to fluorescence confocal microscopy. ODT enables imaging of an identical sample along with multiple time points as the neurons grow. Repeated imaging using a single neuron sample for a long time without the need for additional resources can considerably simplify the experiments (created with BioRender.com).

Alternatively, optical diffraction tomography (ODT) is a 3D quantitative phase imaging technique that enables label-free volumetric quantitative imaging [36]. ODT reconstructs the refractive index (RI) of a sample. As the intrinsic optical property of the sample serves as an imaging contrast, this technique does not require fixation or labeling procedures [37]. This makes ODT suitable for long-term monitoring of live biological samples and enables to image the sample whenever required during the incubation process (Fig. 1). ODTs have been used for imaging various types of biological samples suspended in solution or cultured on 2D surfaces [38–42]. More recently, they have also been applied to samples with extended thicknesses or cells cultured in 3D environments [43–45].

In this study, we show how ODT can be utilized to continuously monitor 3D neuronal growth in *in vitro* cultures using Matrigel as a 3D scaffold. The label-free neuron imaging capability of ODT without any additional sample preparation, such as fixation or labeling, was demonstrated. The 3D images of neurons recorded using confocal microscopy and ODT were compared to verify the accuracy of the neuronal morphology shown in the RI tomograms. Individual neuronal growth was monitored for 4 days using ODT, and the measured tomograms were used for neuron morphology analysis.

## 2. Materials and methods

### 2.1 Three-dimensional neuronal culture with Matrigel

Neurons were obtained from the cortex of E18 Sprague–Dawley rats (Koatech, South Korea). The cortex was removed from the brain by a microsurgical procedure and rinsed with Hank’s buffer salt solution (HBSS; Welgene, South Korea). After the dissection, the cortical cells were dissociated using a micropipette with HBSS. The dissociated cells were centrifuged at 1000 rpm for 2 min, and the media were replaced by plating media (PM). The PM contained neurobasal media supplemented with B-27 (Invitrogen, USA), 2 mM Glutamax (Gibco, USA), 12.5 μM L-glutamate (Sigma-Aldrich, USA), and 1% penicillin-streptomycin (Gibco).

The cell suspension solution was mixed with a gel solution to create a supporting structure for culturing the neurons in 3D. Matrigel (Basement Membrane Matrix, LDEV-free; Corning, USA) was used as a scaffold. After the addition of Matrigel into the cell suspension solution, the final concentration of the cell suspension was 0.5×10^6^ cells/mL, and the final concentration of Matrigel was 7.5 mg/mL.

A silicone elastomer (Sylgard 184, Dow Corning, USA) mold was created on a confocal dish (SPL Life Science, South Korea) to form circular cavities with a diameter of 2 mm and a height of 200 μm. A total of 0.6 μL of the cell-gel mixture was loaded into each cavity and placed in a humidified incubator (5% CO_2_, 37 °C) for 30 min for gelation of the Matrigel. After the gelation, 2 mL of PM was loaded, and the sample was cultivated in the incubator (5% CO_2_, 37 °C, Forma Series II, Thermo SCIENCIFIC). Every 3 days, half of the media was exchanged with fresh media without L-glutamic acid. During multiple time points imaging, the samples were cultivated in the incubator (5% CO_2_, 37 °C, BB15 stainless steel CO_2_ Incubator, Thermo Scientific). All experiments were performed in accordance with the guidance of the Institutional Animal Care and Use Committee (IACUC) of Korea Advanced Institute of Science and Technology (KAIST), and all experimental protocols were approved by IACUC of KAIST.

### 2.2 Optical diffraction tomography

The cultured neurons in their live states were imaged using ODT for label-free quantitative measurements. A custom-built ODT system based on Mach–Zehnder interferometry was utilized for the measurements. Coherent laser light with a wavelength of 457 nm (Cobolt TwistTM; Cobolt, Sweden) was split into a sample and a reference beam using a beam splitter. The sample beam was rotated around the sample using a digital micromirror device (DMD), which allowed fast and stable control of the illumination angle [46, 47]. Time-multiplexed sinusoidal patterns on the DMD diffracted the laser light at various angles.

While the coherent plane-wave laser light illuminated the sample, the position of the sample stage was controlled by a motorized stage (MLS203-1; Thorlabs, Inc., USA). This allowed automated tomogram measurements in adjacent tiles for an extended lateral field of view. A detailed description of the algorithm utilized for stitching the tiles can be found elsewhere [48]. The transmitted laser light after scattering through the sample was collected by an objective lens (UPlanSAPO20X, numerical aperture (NA) = 0.75; Olympus Inc.), and interference patterns were created with the reference beam. The holograms were then recorded using a CMOS camera (LT425M-WOCG, Lumenera Inc.).

A 3D RI tomogram was reconstructed from 335 holograms generated from all different illumination angles in two steps. The first step was to calculate the 2D amplitude and phase information using the phase retrieval process [49]. The next step used the Fourier diffraction theorem for the reconstruction of a 3D RI tomogram from the 2D field data retrieved in the previous step [50]. The resolution of the system was 0.17 μm in the lateral direction and 1.4 μm in the axial direction. The field of view of a single tomogram obtained by the imaging system was 200 × 200 μm^2^, which required approximately 10 s for the measurement and 17 min for the reconstruction. To compensate for the uncollected scattering information owing to the limited NA, an interactive regularization algorithm based on the non-negativity constraint was used [51]. Details on the principle of ODT and the reconstruction algorithm can be found elsewhere [52, 53].

### 2.3 Fixation and immunostaining protocol for confocal microscopy

Neurons cultured in Matrigel were also imaged using a confocal microscope to compare the neuronal morphology obtained from the confocal microscope and ODT. Cultured neuron samples were fixed in 4% paraformaldehyde solution (Biosesang, South Korea) for 6 h in a refrigerator. The samples were gently rinsed three times with phosphate-buffered saline (PBS). To enhance the permeability of the samples, they were treated with 1% Triton X-100 (Sigma-Aldrich) for 6 h at room temperature (25°C) and rinsed three times with PBS. The samples were then treated with the blocking solution, 6% bovine serum albumin solution (BSA, Sigma-Aldrich), overnight at room temperature and rinsed three times with PBS. The samples were incubated at 37 °C overnight in a 1.5% BSA solution with anti-β-III-tubulin (anti-rabbit, 1:500, Sigma-Aldrich) and rinsed three times with PBS. In sequence, the samples were treated with a secondary antibody. The sample was treated with Alexa Fluor 488 (anti-rabbit, 1:500, Invitrogen) with 1.5% BSA solution for 6 h in an incubator (37 °C) and rinsed with PBS three times. PBS with Hoechst 33342 (2.5 μg/mL, Sigma-Aldrich) was added to the samples for 2 h at room temperature and was rinsed three times with PBS. The confocal microscope (LSM 880, Carl Zeiss, Germany) utilized for imaging had a water immersion objective lens (40×, NA = 1.2, C-Apochromat 40x/1.2 water immersion objective, Carl Zeiss, Germany).

### 2.4 Soma and neurite segmentation

A raw 3D RI tomogram of a cultured neuron sample includes uneven backgrounds, which require segmentation of the soma and neurites of each neuron for further analysis. The soma of a neuron was segmented based on RI values. It was identified by voxels with RI values higher than 1.3424. To segment the neurites, a strategy different from the RI thresholding was utilized. The Simple Neurite Tracer tool [54], which is part of the plugin package in Fiji, was utilized [55]. The segmentation process using the Simple Neurite Tracer involved the following steps. First, the starting point of a neurite was selected by a trained user, and the next point further away from the starting point was selected by the user. The tool automatically connected the two points in 3D based on the highest RI values in the path. This was repeated until the connection reached the end of the neurite. When all neurites were segmented using the Simple Neurite Tracer tool, the data were subjected to Hessian-based analysis, which improved the accuracy of the selected path.

### 2.5 Neurite growth direction analysis

The segmented neurites were further analyzed to calculate the initial growth directions. After the 3D volume of each neurite was segmented by the Simple Neurite Tracer tool, the skeleton of each neurite was found by the “bwskel” function in MATLAB(The Mathworks Inc., USA). A low-pass filter and averaging five consecutive pixel positions were then applied to the skeleton to obtain a continuous path for the skeleton. The direction vectors of the skeleton were calculated based on the relative position differences between two consecutive pixels and normalized to the length of a pixel. The growth direction vectors of the secondary neurites growing from the other neurites were considered independently.

## 3. Results and discussion

### 3.1 Refractive index tomogram of neurons cultured in 3D

Neurons cultured inside Matrigel were measured using ODT for the label-free recording of live neuron growth in a 3D environment (Fig. 2; see Methods). An ODT optical system was custom-built to image the neurons (Fig. 2A). The laser beam illuminating the sample rotated around the sample at 335 different angles to generate 335 holograms. The holograms were used to retrieve both the amplitude and phase distributions of the samples (Fig. 2B). A single 3D RI tomogram was reconstructed from the measured multiple 2D optical field images [56]. Multiple tomograms were often stitched together to achieve a larger field of view for measuring the entire structure of the neurons. The reconstructed RI tomogram of the neurons revealed the 3D morphology of the neurons in their live state (Fig. 2C). The region of interest (ROI) in Fig. 2C shows a single neuron cultured for 2 days *in vitro* (DIV). Multiple 2D lateral image slices of the tomogram showed a continuously varying spatial distribution of the neurites growing out from the soma. The 3D RI-based-rendered data of identical neurons are shown in Fig. 2D. Neurites of the neurons had RI values below 1.3424. The somata of the neurons had higher RI values, which indicates a higher molecular concentration in the volume. The measured RI tomograms clearly demonstrate the potential of the ODT to visualize the 3D structure of live neurons cultured in Matrigel.

**Fig. 2.**
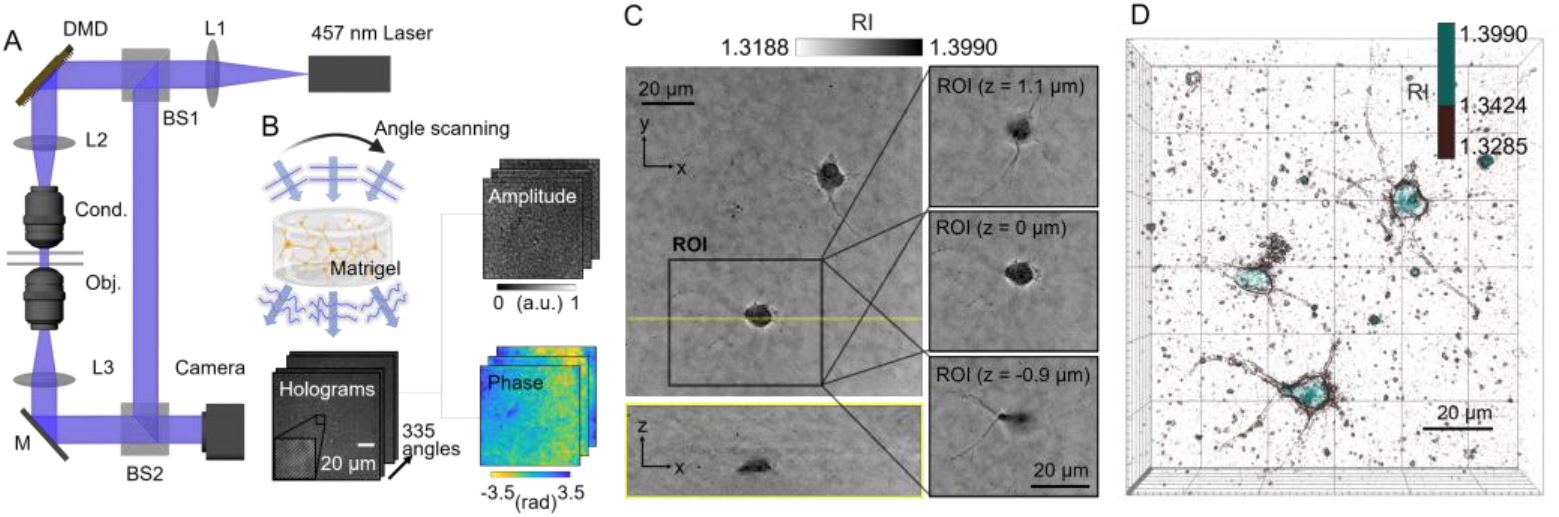
3D RI measurement of cultured neurons using ODT. **A** Schematic of the ODT system based on the Mach–Zehnder interferometry with a DMD. **B** The neurons were cultured inside a Matrigel scaffold and imaged in 3D using ODT. The laser beam illuminated the sample from 335 different angles and each illumination generated a hologram. From the 335 holograms, the amplitude and the phase information of the sample were retrieved, from which the 3D RI tomogram was reconstructed. **C** Representative tomogram of a cultured neuron in a Matrigel scaffold. The ROI indicates a neuron and its 2D sections at various axial positions. **D** RI-based 3D-rendered volume of the tomogram shown in Fig. 2C. The measured RI values were higher (1.3424–1.3990) in the soma than those in the neurites (1.3285–1.3424).

### 3.2 RI tomogram and confocal fluorescence with compatible neuron morphology

To verify that the morphology of neurons measured with ODT is compatible with conventional confocal fluorescence microscopy, the morphology data of the neurons obtained by two different imaging modalities were compared (Fig. 3). After the cortical neurons from E18 rats were cultured in Matrigel (see Methods), their RI tomograms were measured by ODT after 3 DIV. Because confocal microscopy requires external fluorescence labels for imaging, the neurons were fixed immediately after the ODT measurement for immunostaining. The fixed neurons were imaged using the ODT again before the immunostaining procedure, followed by confocal microscopy after staining.

**Fig. 3.**
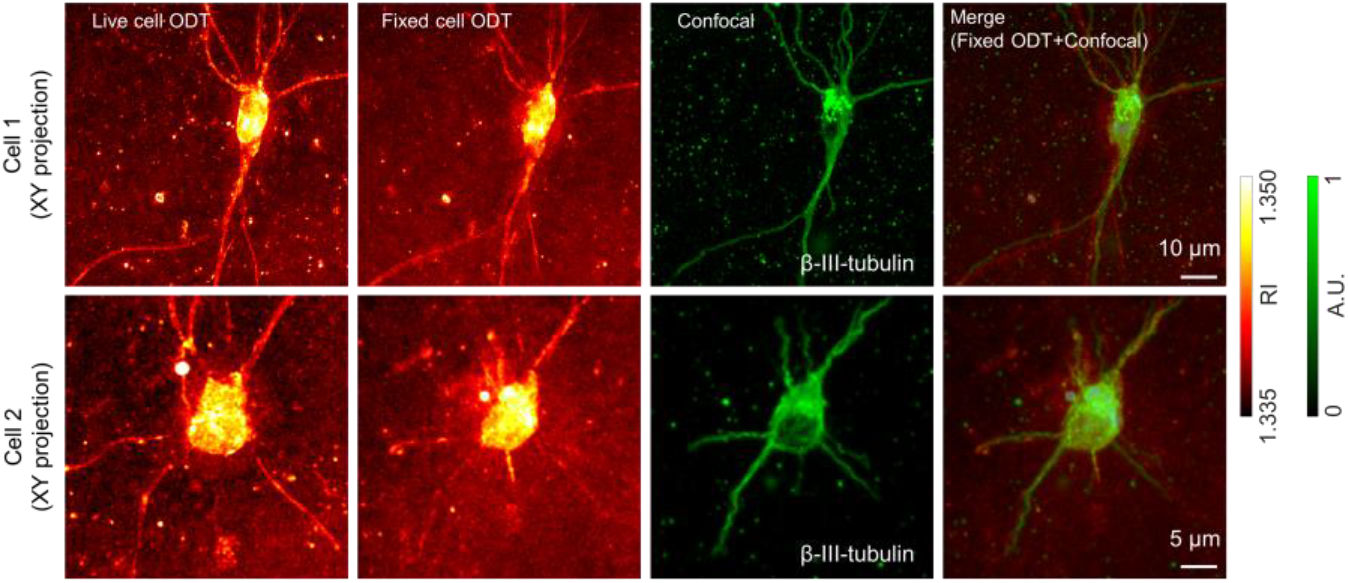
Comparison between the XY maximum intensity projection images of the 3D data obtained by ODT and confocal microscopy. Images of two different neurons after 3 DIV were obtained. Neurons at both live and fixed states were imaged using ODT. After fixation, the β-III-tubulins of the neurons were labeled with fluorescence via immunostaining and imaged by a confocal microscope. The morphology of the fixed neuron also correlated well in the images measured by ODT and confocal microscopy (merged images).

When the RI tomograms of the neurons were compared before and after the fixation procedure, the 3D morphology of the neurons did not seem to be significantly altered by the fixation. However, the RI values in the background were slightly elevated, and some of the original high-RI components in the neurites were eliminated during fixation. A similar effect of the fixation procedure on the RI of cells has been previously reported [57, 58]. Although the effect was highly dependent on the cell type, the RI values and contrast between the subcellular components decreased. When the RI tomogram and confocal fluorescence data were compared, the structure of the fixed neurons did not vary, except for some mismatches due to the different imaging modalities. Thus, it was confirmed that the morphology of neurons measured by ODT tomogram was identical to that from conventional imaging methods. Furthermore, the neurons in their live state showed an ODT tomogram with better image quality than at their fixed state. This result indicates that ODT can considerably simplify the imaging procedure and reduce the resources required for the experiments. ODT can show compatible morphological data as the confocal microscope with minimal disturbance of the live neurons.

### 3.3 Segmentation of 3D RI tomograms for neuron morphology analysis

For systematic analysis of individual neurons, the soma and neurites of the neurons were segmented individually (Fig. 4A). First, the soma of a neuron was successfully segmented from the tomogram by selecting the voxels with RI values higher than 1.3424. This thresholding value yielded the best segmentation results for the soma, and the debris in the background was excluded. Next, neurites were segmented using a plugin tool in ImageJ called Simple Neurite Tracer. Unlike segmenting the soma, the simple one-step segmentation based on RI thresholding did not work for neurites. This was because the RI ranges of the high-RI background noises and the neurites often overlapped each other. The spatial context was required to effectively segment the neurites. After both the soma and neurites were segmented, the two masks were added together to create an entire 3D mask of a neuron. By applying the mask to the original RI tomogram, clean data showing only neurons were obtained. Identical data were rendered in 3D (Fig. 4B).

**Fig. 4.**
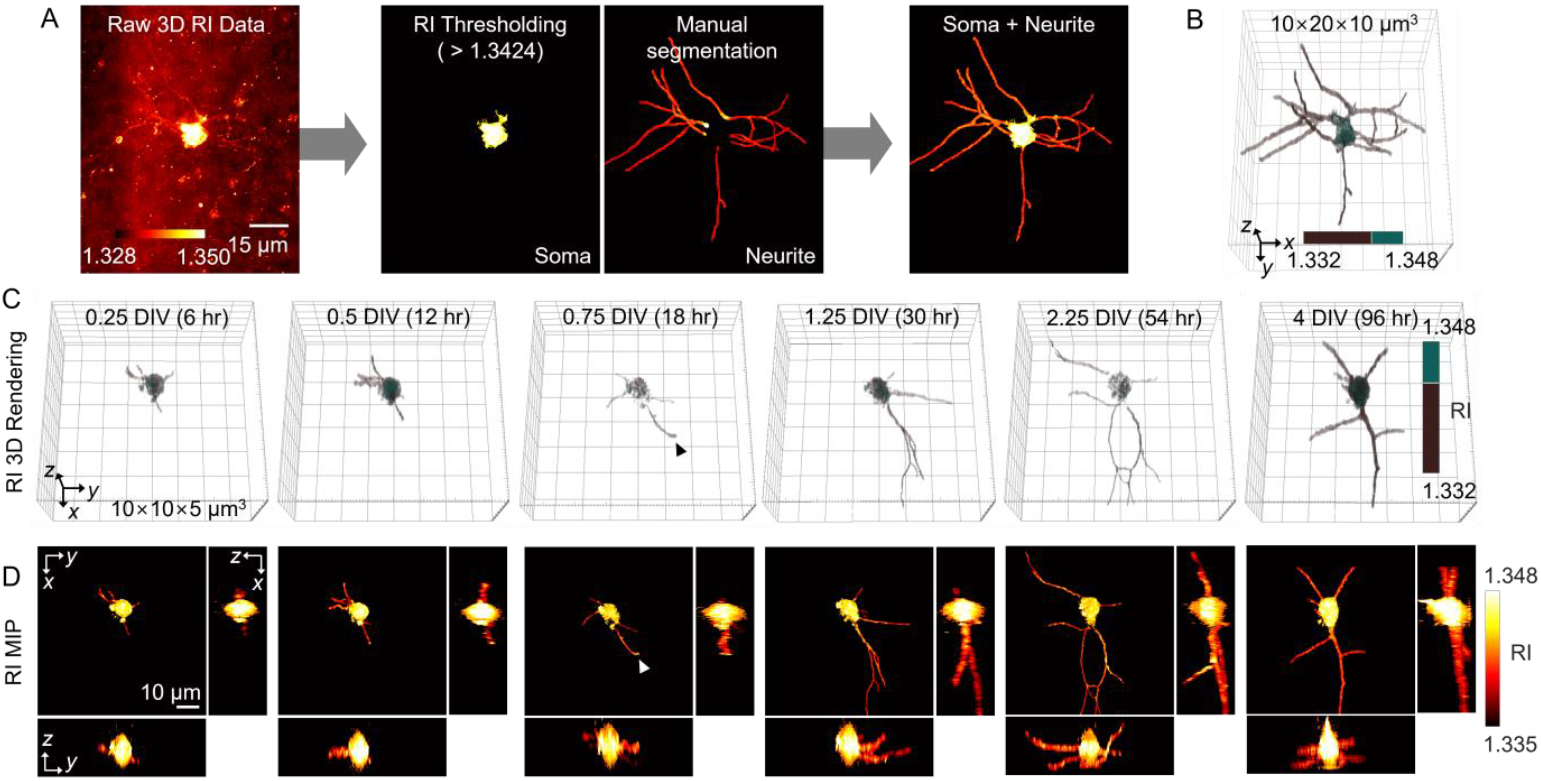
3D morphology measurements of neuron growth over time using ODT. **A** Raw RI tomograms were preprocessed to segment the soma and neurites for neuronal morphology analysis. **B** 3D rendered RI tomogram of a neuron after the segmentation procedure for both the soma and the neurites. **C** Time-lapse RI tomograms obtained by tracking a single neuron growth for 4 days. The tomograms shown here were obtained after soma and neurite segmentation. **D** MIP images of the 3D RI tomograms in Fig. 4C along three different planes.

The major advantage of imaging neurons using ODT is the ability to visualize neuronal morphology at the single-cell level without interrupting neuron growth. The 3D structures of individual neurons were measured by ODT for 4 days, starting from when they were seeded into Matrigel. The soma and neurites of each neuron were segmented from the raw tomograms. Tomograms were obtained every 6–12 h during culture. The results were then rendered in 3D to visualize the growth of neurons inside a 3D Matrigel (Fig. 4C). Each tomogram was used to generate maximum intensity projection (MIP) images in three different axial directions (Fig. 4D). They demonstrated the 3D neurite formation procedure. Minor processes developed at 0.25 DIV(6 hours) and a major neurite (‘putative axon’) outgrowth were observed at 0.75 DIV(18 hours) with a slightly elevated RI at the end.

### 3.4 Neurite growth direction with asymmetric polar angle distribution

To further demonstrate the capability of 3D label-free imaging of individual neurons cultured in 3D, the growth directions of the neurites were investigated. Individual neurites in each neuron were segmented from the measured tomogram, and their distribution in the 3D space was analyzed (Fig. 5). The MIP images in the lateral plane (*xy* plane) were modified into the axial position colormaps, where the color of each pixel was determined by its axial (*z*-axis) position. Once the neurites were segmented from the original tomogram, their skeleton was calculated, and the initial growth direction of each skeleton was obtained (see Methods).

**Fig. 5.**
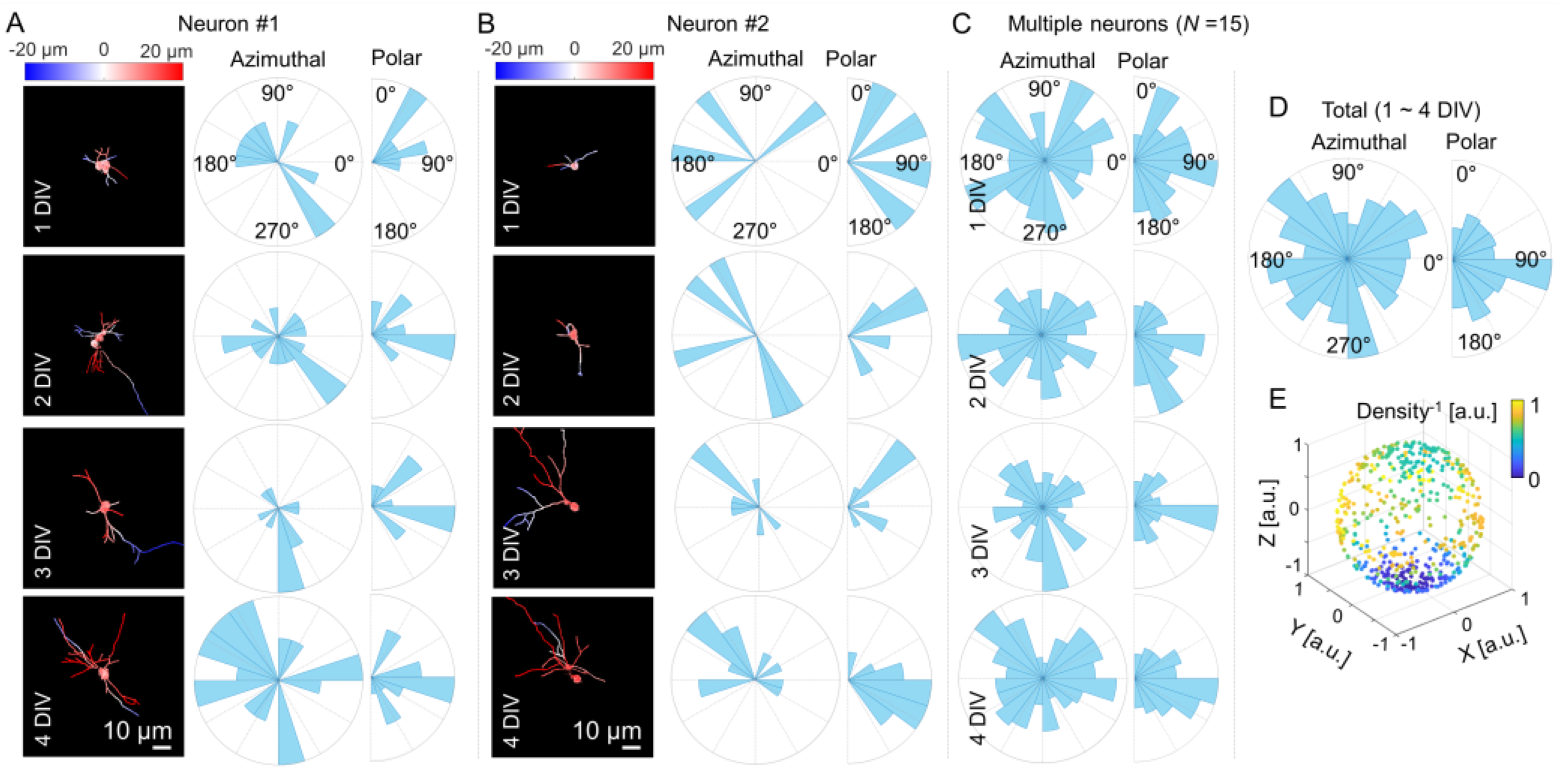
3D growth analysis of individual neurites over time using ODT. **A, B** Each section shows the neurite growth profile of a single neuron along time. Time-lapse tomograms of single neurons were preprocessed and projected onto the *xy* plane. Axial position colormap represents the axial (*z*-axis) position of the pixel by color. **C** Neurite direction histogram for multiple neurons (*n* = 15) over time. **D** DIV-averaged direction histogram. **E** Neurite growth directions are shown in a 3D space when the soma is located at the origin. The color of each data point describes the average distance of the closest 40% of all data points, which is inversely proportional to the local density at the position.

The 3D morphology of two different neurons from one to four DIV is shown in Figs. 5A and 5B. The directions of individual neurite growth are represented as angular histograms in the spherical coordinate system showing the distribution in the azimuthal and polar angles. The neurite direction data of multiple (*n* = 15) neurons are presented together to show the overall behaviors (Figs. 5C–D). As neurons grew, neurites grew to have a more uniform distribution in the azimuthal angle. However, in the polar axis, the neurites tended to grow more toward the lower hemisphere. For the better visualization of the growth direction of neurites, each directional vector was projected onto a sphere, and each point was assigned a color based on the local density near the data point (Fig. 5E). The density was the highest in the lower hemisphere.

## 4. Conclusion

This study demonstrates the proof of concept for label-free monitoring of 3D-cultured neurons using ODT. Primary cortical neurons cultured in Matrigel were used as samples, but the imaging target can be readily extended into a co-culture of neurons and astrocytes or other types of 3D cultures. Moreover, several advanced fluorescent labels that can be used for imaging cells in their live state have been developed. For example, NeuO has been developed to selectively label live neurons, although the staining remained stable for 36 h [59]. CDr10b was also developed to label live microglia cells, and the signal was robust for more than 18 h [60]. Although fluorescent labels can be modified or utilized for live-cell imaging, RI is the physical property of the sample that is not affected by the imaging frequency or intensity of illumination light. For this reason, ODT can be considerably advantageous compared to fluorescence microscopy when tracking a single sample is desirable for studying the developmental processes of neuronal growth. However, the molecular or cellular specificity information provided by fluorescence microscopy can be combined with the RI tomograms because the ODT system cannot provide the molecular or cellular specificity by itself. Therefore, in addition to obtaining RI tomograms using ODT, measuring fluorescence images to provide cellular-level selectivity with the use of live-cell fluorescent dyes, such as NeuO and CDr10b, would provide even more affluent information about the sample over time. Furthermore, there have been recent advances in predicting additional information about biological samples using deep learning [61–63]. By extending these approaches into 3D neurons, the information conventionally obtained by labeling can be acquired from time-lapse label-free RI tomograms.

In conclusion, our study suggests that ODT is a unique tool for monitoring neuronal growth in 3D *in vitro* cultures by measuring the morphology of neurons without fixation or labeling procedures. Imaging the morphology of live primary neurons in 3D can provide fundamental information about the most basic building components of the mammalian central nervous system. Our method can also be applied to other types of 3D cultures using different cell types or culture protocols. Continuous monitoring of individual samples for an extended period will provide an invaluable dataset for neuroscience and drug discovery. Long-term monitoring data obtained by ODT would considerably benefit experiments such as testing drugs, studying disease progression in various disease models, and investigating the regeneration process after injury.

## Data availability

Data underlying the results presented in this paper are not publicly available at this time but may be obtained from the authors upon reasonable request.

